# Bounded Multiplicative Dynamics Govern Axonal Conduction Slowdown

**DOI:** 10.64898/2026.03.03.709326

**Authors:** Shimon Marom

## Abstract

Conduction velocity slows along axons, yet the terminal-to-initial ratio *ρ* = *v*_end_*/v*_start_ is branch-length invariant and right-skewed. A bounded multiplicative framework is introduced, in which local geometric and kinetic factors compound proportionally, but only within a finite distal domain set by termination conditions that saturate the effective multiplicative depth. The model accounts for the observed stability of the slowdown distribution across lengths and yields discriminating experimental signatures that distinguish between structural load and kinetic reserve depletion.

## INTRODUCTION

Under most physiological activity regimes, including spontaneous network activity and low-to-moderate stimulation rates, axons function as robust conducting structures, reliably transmitting action potentials across distances and levels of axonal branching [1–7]. This robustness does not imply trivial dynamics. Axonal propagation is shaped by geometry, membrane properties, and boundary conditions, and classical cable-theoretic analyses show that conduction velocity is sensitive to axial load, diameter changes, and sealed-end terminations [8–10]. These effects are generally interpreted as modulations of a fundamentally reliable signaling process, rather than as transformations of representational content conveyed by the spike itself.

Experimental and computational studies have long noted that axonal inhomogeneities can induce progressive slowing as spikes propagate away from their site of initiation [11]. The slowdown is not merely transient or tied to particular structural contingencies, but instead exhibits a stable statistical organization when examined across large axonal ensembles [12]. As demonstrated in Figure 1, individual velocity profiles display a gradual, predominantly monotonic deceleration along branches. The ratio between terminal and initial velocities, *ρ* = *v*_end_*/v*_start_, is right-skewed and well described by a log-normal distribution, with a typical slowdown of about 30%. Within the measured range (0.6–4.6 mm), there is no statistical evidence for length-dependence of *ρ*, nor for broadening of ln *v*.

**FIG. 1.**
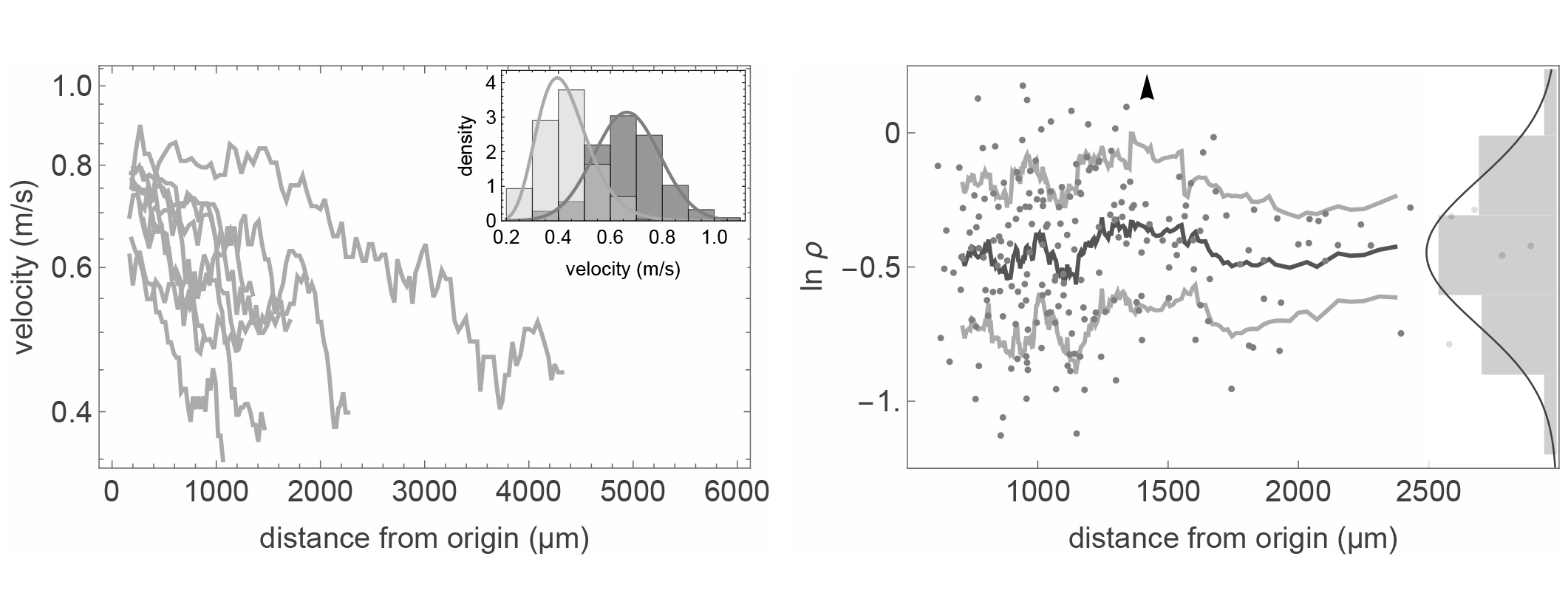
Statistical organization of spike propagation along axons. [adapted from Cohen and Marom [12]]. *Left*: Representative log–linear conduction-velocity profiles for axons of different lengths. *Inset (left)*: Histograms of *v*_start_ (dark gray) and *v*_end_ (light gray) for branches (*n* = 214) selected to originate near the somatic spike-initiation zone by requiring an initial time-of-arrival (TOA) of at most 200 *µ*s relative to the triggering somatic spike (corresponding to an origin within ∼ 130 *µ*m given 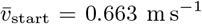). *Right* : Scatterplot of ln(*v*) versus branch length for the same *n* = 214 subset, where *ρ* = *v*_end_*/v*_start_ denotes the slowdown ratio (plotted up to 3 mm for visual clarity), overlaid with a rolling mean and rolling standard-deviation bands (ρ1 SD of ln(*v*); window size = 20). One point lies above the displayed *y*-range; it is retained in all analyses and indicated by an arrow. Slowdown shows no systematic dependence on length (Pearson *r* = 0.018, *p* = 0.71; Spearman *r*_*s*_ = 0.044, *p* = 0.36), and dispersion shows no evidence of length-dependent broadening (Breusch–Pagan test on residuals from a linear regression of ln(*ρ*) on length: *p* ≈ 0.56). *Inset (right)*: Distribution of ln(*ρ*) with a normal fit (*µ* = −0.44, *σ* = 0.27).

These observations indicate that propagation reflects the compounded effect of many small, local influences, even as the effective multiplicative depth remains bounded. Several mechanistic scenarios could instantiate this distributed compounding, including spatial gradients in channel density or other structured inhomogeneities. A minimal model is introduced, grounded in cable-theoretic excitability, in which distal boundary constraints impose a terminal multiplicative pre-factor that is a function of distance to the tip. This boundary-anchored contribution dominates the net effect of the multiplicative updates and yields slowdown statistics that are effectively independent of total trajectory length. Accordingly, the distal contribution is treated as an effective deterministic termination operator that confines the multiplicative contribution governing *ρ* to a finite distal spatial domain near the sealed end, thereby preserving the observed length-invariance of Var[ln *v*].

The proposed model resembles transport toward a distal boundary condition: regardless of upstream distance, approaching the boundary imposes a boundary-set deceleration. Any axonal terminal realization that reduces the local safety factor (conduction margin) over a finite coupling depth can limit propagation via structural, kinetic, or mixed mechanisms [13], including involvement of intra-axonal organelles (e.g. [14]). The formulation proposed here yields realization-informative signatures because distinct distal mechanisms differentially modulate both perturbation-induced changes in 𝔼 [ln *ρ*] and the effective multiplicative depth.

## MULTIPLICATIVE SLOWDOWN AND THE FAILURE OF UNBOUNDED ACCUMULATION

The compounded influences are formalized by expressing conduction velocity along a branch as a recursive product of local multiplicative factors. Let *v*_*n*_ denote the velocity in segment *n*, and let *κ*_*n*_ ∈ [0, *∞*) be a dimensionless factor representing the local geometric or kinetic modulation of velocity over that increment. Then *v*_*n*+1_ = *κ*_*n*_ *v*_*n*_, so that over *N* segments the terminal velocity satisfies

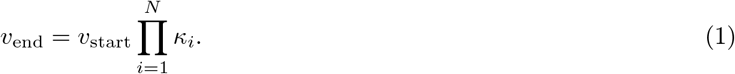

“Segment” denotes a coarse-grained spatial increment over which local proportional modulation of the source-to-load balance is summarized by a single multiplicative factor *κ*_*i*_. It is not intended as an anatomical or electrotonic unit, and its scale is a modeling resolution parameter.

Formally, this is a one-dimensional multiplicative propagation scheme, in which transport along a path is described as a fixed reference scale multiplied by a product of local factors that represent successive inhomogeneities; local geometric and kinetic influences that rescale spike conduction velocity along the axon. Writing the end-to-start ratio as *ρ* = *v*_end_*/v*_start_ gives

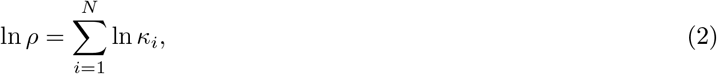

consistent with a right-skewed, approximately log-normal distribution for *ρ*. This log-normality of *ρ* is assessed directly from the near-normality of ln *ρ* (Figure 1, right panel) [12]; it is not (and in fact, cannot be) inferred from the marginal forms of *ρ*_start_ and *ρ*_end_.

Note that the above multiplicative scheme entails a correlation of the first and second moments of ln *ρ* with travel distance. This, however, does not align with empirical observations (Figure 1, right panel). Thus, local geometric and biophysical influences might still modulate propagation proportionally, such that *ρ* = Π _*i*_ *κ*_*i*_, compounding along the trajectory; but the overall variability of slowdown remains constrained. This constrained multiplicative structure provides the basis for the formal treatment offered here.

## BOUNDARY-ANCHORED COARSE GRAINING

Conceptually, *v*_start_ serves as a reference scale relative to which multiplicative modulation operates. A pertinent question, therefore, is whether *v*_start_ can be treated as approximately constant. For a given neuron, *v*_start_ appears sufficiently stable to serve in this role, even if it varies across neurons and preparations. This is consistent with the Hodgkin–Huxley view of local excitability as a stable dynamical property [15], and with empirical observations of stable spike amplitude and waveform near the initiation site. Accordingly, the relevant stability is not constancy across neurons, but stability within each neuron across its branches.

Thus, axonal initiation and termination impose fixed constraints: conduction begins near a stable somatic initiation zone and ends at a distal termination, typically at a tapering tip. Classical cable theory shows that sealed-end terminations, together with geometric tapering toward the tip, influence voltage and current over approximately one electrotonic length (λ) from the distal end, whereas more proximal (i.e., upstream) regions remain comparatively insensitive to the termination in the subthreshold cable sense [8]. These distal impedance effects redistribute axial current and reshape local voltage gradients near the tip. In special cases they can yield local acceleration [9]. However, as discussed below, in the present regime their net effect is a reduced safety factor and an overall slowdown, consistent with the observed phenomena, when regenerative inward current becomes marginal. (Note: The classical λ bound pertains to linear subthreshold perturbations, whereas conduction velocity is governed by a distributed, state-dependent source-to-load balance in which the driving voltage at the wavefront must overcome the effective downstream impedance, including both axial resistance and capacitive loading. Thus, “distal sensing” in the regime treated here should be understood as a boundary condition on active propagation, not as long-range electrotonic influence.)

In principle, conduction along a sufficiently extended axon could reflect contributions from two spatial domains. Far from a sealed or tapering termination, velocity could remain approximately constant or drift gradually in a way that need not be strictly multiplicative. If such an upstream contribution accumulated with distance, one would expect the terminal slowdown ratio *ρ* = *v*_end_*/v*_start_ to depend systematically on branch length. Yet, at least across the five-fold range of lengths shown in Figure 1 (right), the mean of ln *ρ* remains effectively invariant, and no gross broadening of the slowdown distribution is evident in the aggregate data. Absent fine-tuned cancellation, the most economical interpretation is that any proximal changes in velocity do not contribute appreciably to the statistics of *ρ*, and can be absorbed into the branch-specific reference scale *v*_start_. These considerations motivate restricting the present analysis to a bounded distal contribution near an effective termination, and the toy model below focuses on the corresponding finite multiplicative depth within the distal coupling zone set by the terminal boundary.

Let *η*(Λ) denote a deterministic factor representing the net distal constraint. It approaches unity when the constraint is weak and decreases as termination becomes more restrictive. Here, Λ (capital lambda) denotes an effective *active* range over which distal constraints can influence regenerative propagation, to distinguish it from the passive electrotonic length λ. The range Λ may exceed λ, which delimits linear subthreshold locality.

Importantly, *η*(Λ) is defined at the level of an effective realization rather than a microscopic mechanism. It may arise from load or impedance effects, including sealed-end termination and taper-driven current focusing; from excitability effects, including graded distal reductions in sodium conductance or channel availability, increased outward currents, or other losses of safety factor; or from combinations of these. In this sense, *η*(Λ) summarizes the distal renormalization of the local source-to-load balance that ultimately sets *v*_end_.

The normalized end-to-start ratio can then be written schematically as

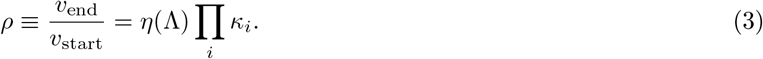

This ratio is dimensionless by definition. Hence, *η*(Λ) denotes the coarse-grained deterministic bias imposed by termination geometry and the associated boundary-coupled excitability reserve, whereas the *κ*_*i*_ encode residual spatial heterogeneity within the boundary-coupled distal segment around that bias. This separation is a modeling convention that prevents the boundary contribution from being absorbed into the empirical mean of ln *κ*_*i*_.

Also, for later use, it is helpful to decompose the effective distal bias into two conceptually distinct components,

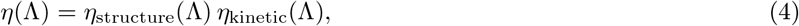

where *η*_structure_ summarizes effectively time-independent distal constraints on the source-to-load balance [8–10, 13], including sealed-end termination, tapering, and other structural or nearly-static spatial factors (for example diameter variation, branching, axial resistivity changes, or fixed gradients in channel expression). In contrast, *η*_kinetic_ summarizes history- and voltage-dependent adaptive processes that reduce excitability reserve [6, 10, 16–19], including changes in channel availability, inactivation- and accommodation-like losses, and other kinetic reductions in regenerative margin (see Supplementary Figure S1 for a Hodgkin-Huxley-based illustrative proxy and a waveform signature of reduced availability).

Eq. 4 is a log-additive bookkeeping device, not an assertion that structural and kinetic contributions are mechanistically independent. Any non-separable coupling can be absorbed into the Λ-dependence of the two factors, or represented explicitly by an interaction term. The decomposition is not claimed to be unique; it is chosen to keep the model implementation-agnostic while enabling predictions that discriminate among realization classes. It is a phenomenological partition of a deterministic distal-bias operator that shifts the slowdown mean, while the dispersion plateau remains governed by the bounded multiplicative depth. Empirically, under typical conditions *η*(Λ) *<* 1, so the distal boundary acts as a net decelerating bias that aggregates load/impedance and excitability-reserve effects into a single renormalizing operator.

## BOUNDED MULTIPLICATIVE DEPTH AND LENGTH-INVARIANT DISPERSION

A simple analytic construction clarifies how the distal factor *η*(Λ) can preserve the log-normal form of slowdown while suppressing the length dependence of Var[ln *ρ*] characteristic of a multiplicative propagation scheme. Let

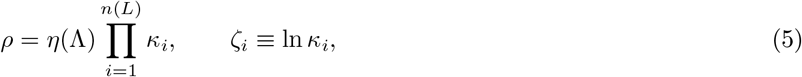

where *L* denotes distance measured *from the sealed termination back toward the soma*, and *ζ*_*i*_ are local log-increments with mean *µ* and variance *σ*^2^. Introducing *ζ*_*i*_ linearizes multiplicative compounding, so that ln *ρ* can be analyzed using standard results for sums of weakly dependent increments.

In a passive one-dimensional conductor, the number of effectively independent increments *n*(*L*) grows linearly with distance, yielding

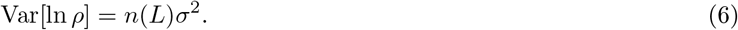

This is the source of length-dependent variance in multiplicative one-dimensional propagation schemes. In an axon, however, the sealed or tapering termination imposes a boundary condition whose influence decays with distance from the tip with a characteristic scale Λ. Beyond that scale, additional distal cable couples only weakly to the terminal ratio *ρ* through the distal boundary condition, so the effective number of statistically independent multiplicative contributions that can influence the terminal slowdown saturates rather than growing linearly. This saturation concerns sensitivity of the *terminal ratio* statistic *ρ* = *v*_end_*/v*_start_ to boundary-coupled fluctuations; upstream heterogeneity may still contribute to absolute travel time without remaining coupled to the sealed-end source-to-load margin that determines *v*_end_.

A minimal coarse-grained representation for this boundary-limited coupling is a saturating effective depth,

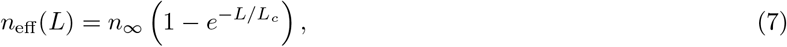

where *L*_*c*_ is an effective coupling length Λ (not necessarily identical to the passive electrotonic λ) and *n*_*∞*_ is the *saturated depth*. This formulation does not assume that local multiplicative factors differ intrinsically along the axon. Rather, it reflects that only fluctuations that remain coupled to the distal boundary can influence the terminal ratio, and this influence is strongest when propagation is near threshold, where small variations in local conductances or impedance produce disproportionately large changes in regenerative balance. The saturation of *n*_eff_ is therefore a statement about coupling and sensitivity, not about the absence of heterogeneity elsewhere. The resulting slowdown can be written as

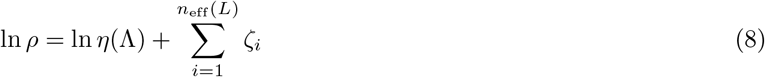

(Here *n*_eff_(*L*), though continuous, is used in a coarse-grained sense). Thus,

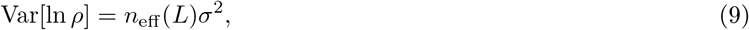

And

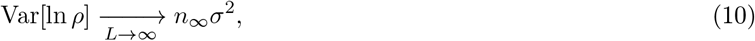

a finite constant, while the mean slowdown continues to depend on ln *η*(Λ) and the saturated value of *n*_eff_. This toy model shows that saturation of the effective multiplicative depth *n*_eff_(*L*) suppresses the growth of Var[ln *ρ*] by limiting the number of independent multiplicative perturbations that remain coupled to the terminal statistic, whereas deterministic boundary factors embedded in *η*(Λ) (including the components *η*_structure_ and *η*_kinetic_) primarily shift 𝔼 [ln *ρ*] through ln *η*(Λ) without setting the dispersion plateau. The result is a bounded multiplicative process with log-normal form and length-invariant dispersion, consistent with the empirical slowdown observed in axons.

A Monte Carlo comparison illustrates that removing saturation in *n*_eff_(*L*) produces length-dependent broadening of Var[ln *ρ*] over the measured range, whereas a bounded effective depth reproduces the observed length-invariant dispersion (Fig. 2).

**FIG. 2.**
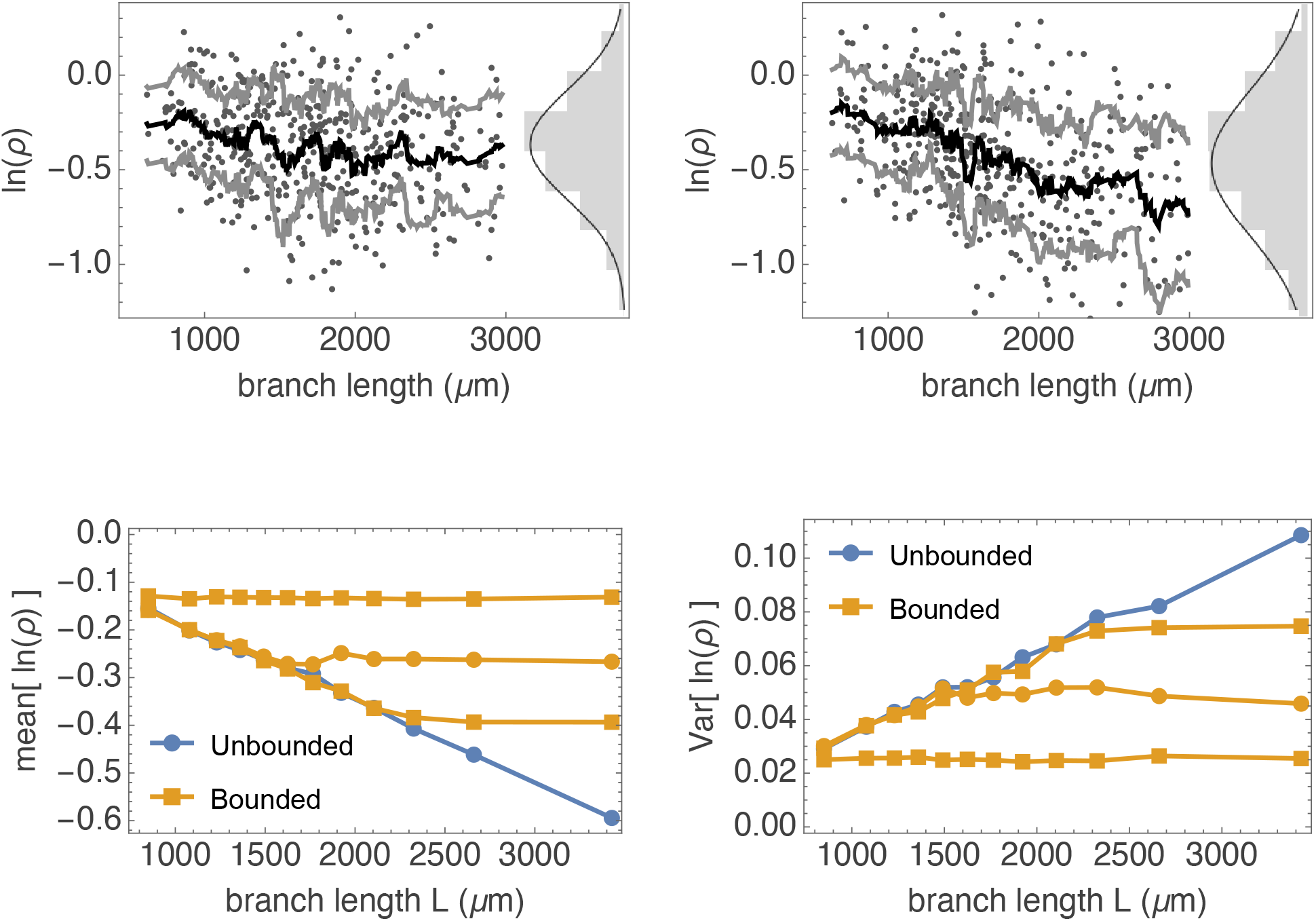
Monte Carlo comparison of bounded and unbounded multiplicative slowdown. Synthetic slowdown ratios *ρ* = *v*_end_*/v*_start_ are simulated under two multiplicative models that differ only in the effective number of compounding steps along a branch: unbounded depth increases with branch length, whereas bounded depth saturates beyond a finite scale (*l*_*s*_). **Top:** Simulated ln *ρ* versus total soma-to-terminal branch length for the bounded (left, *l*_*s*_ = 1600 *µ*m) and unbounded (right) models, with binned central trends and marginal histograms of ln *ρ*. **Bottom:** Length-binned estimates of 𝔼 [ln *ρ*] (left) and Var[ln *ρ*] (right). In the bounded model, both the mean and variance approach a plateau that depends on the value of the saturation depth (*l*_*s*_ = {800, 1200, 2400} *µ*m); in the unbounded model, they grow with length. Simulation details and parameter calibration are given in Supplementary Methods.

## INTERPRETATION AND DISCRIMINATING PREDICTIONS

Products of random factors arise broadly in transport through heterogeneous media, growth processes, and cascade models. In unbounded multiplicative accumulation, the dispersion of ln *ρ* is expected to grow with multiplicative depth. Here, a living excitable medium is analyzed, axons in which *ρ* = *v*_end_*/v*_start_ is approximately log-normal, yet the dispersion of ln *ρ* shows no evidence of broadening over a five-fold range of axonal lengths. This combination of multiplicative form and bounded dispersion constrains the admissible coarse-grained dynamical models, pointing to mechanisms that saturate the effective multiplicative depth rather than allowing unbounded accumulation.

The main claim is that axonal slowdown is captured, at the level of distributional structure, by a bounded multiplicative process modulated by distal boundary conditions:

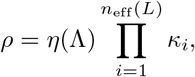

with *η*(Λ) = *η*_structure_(Λ) *η*_kinetic_(Λ). The log-normality and length-invariance of ρ are empirical invariances; the bounded multiplicative form is proposed as a minimal arrangement that satisfies them simultaneously. The argument therefore runs from invariances to a constrained class of mechanisms, not from a distributional fit to a specific microscopic cause. In this sense, identifying individual *κ*_*i*_ with specific biophysical entities, or deriving a unique expression for *η*(Λ), would be underdetermined by the data and would add pseudo-precision rather than insight. Likewise, biophysical simulations that instantiate particular choices of the *κ*_*i*_ and of *η*(Λ) are possible, but they would be illustrative rather than discriminating: many distinct microscopic implementations can realize the same bounded multiplicative law.

At this resolution, the aim is not to enumerate microscopic contributors, but to distinguish distal boundary regimes and their experimentally visible signatures. This motivates three experimentally accessible predictions designed to discriminate between structural load/impedance and kinetic reserve depletion.

*First*, if orthodromic and antidromic spikes are initiated simultaneously along the same branch, their approach to collision creates a transient *effective termination zone* in which the wavefronts annihilate. Within the decomposition *η*(Λ) = *η*_structure_(Λ) *η*_kinetic_(Λ), this paradigm primarily probes *η*_kinetic_: as the fronts converge, regenerative margin should be transiently reduced by availability-dependent kinetics (for example, inactivation- and accommodation-like depletion of available excitatory sodium conductance), yielding measurable slowing relative to unidirectional propagation under otherwise identical conditions. Detection of collision-associated slowing, with minimal manipulation of impedance, would therefore support a non-negligible contribution from *η*_kinetic_(Λ). Conversely, the absence of slowing would point to a predominantly structural contribution, *η*_structure_(Λ), anchored to terminal architecture rather than collision-induced reserve depletion.

*Second*, the mean of the ln *ρ* distribution should increase when spikes are initiated at the far end of the axon and propagate to the soma. Here *ρ* is defined, as above, by *ρ* = *v*_end_*/v*_start_, with “start” denoting the distal initiation site and “end” the soma. This paradigm most directly interrogates *η*_structure_: the large proximal sink and high channel density act as an “open” boundary with *η*(Λ) *>* 1, producing the mirror image of the distal slowing bias in the mean of ln *ρ*. A variant would be to induce sub-threshold depolarization at the soma, thereby probing for an additional contribution from *η*_kinetic_(Λ).

*Third*, for branches whose physical length substantially exceeds the effective distal coupling length *L*_*c*_ (measured from the sealed end toward the soma), the velocity profile is predicted to contain an extended proximal segment with approximately constant, or only weakly declining, velocity, followed by a pronounced distal deceleration confined to a terminal region of order *L*_*c*_. In very long axons, such a “free-running” initial segment should therefore be observable in length–velocity profiles, with the bounded multiplicative slowdown emerging only as the spike enters the sealed-end zone. Because the electrode field in the device used in [12] spans only ca. 2 × 4 mm, the dataset does not include axons long enough to evaluate this asymptotic prediction; testing it would require a recording modality capable of probing substantially longer trajectories.

Together, these predictions outline experiments that reveal how boundary conditions shape the multiplicative process governing axonal propagation. They are not merely descriptive; they are constructed to discriminate among distal boundary regimes (load/impedance versus excitability reserve) that can otherwise generate the same bounded slowdown statistics.

Finally, for completeness, bounded statistics do not compel a single mechanistic narrative. Passive, deterministic impedance transformations, branch geometry, and a small number of strong distal discontinuities can all generate an effective terminal bias; within the present framework such effects are naturally absorbed into *η*_structure_(Λ). A remaining confound is preparation-level state variability, which can be tested directly by asking whether ln *ρ* values co-vary across different trajectories recorded in the same preparation (common-mode covariance), and by controlled global perturbations.

At the broader level, the results place axonal conduction slowdown within a class of self-normalizing physiological processes, in which local variability is preserved but prevented from diverging globally. Earlier work has shown that axonal excitability can be stabilized against structural and kinetic heterogeneity through activity-dependent normalization mechanisms [16–18]. The present findings extend this logic: local geometric and kinetic influences are consistent with proportional compounding along the trajectory, while boundary conditions constrain the net effect observed at the branch terminus. Robust conduction thus emerges not from uniformity, but from constrained variability: a relational balance between local heterogeneity and global boundary conditions.

## SUPPLEMENTARY FIGURE

**Figure S1.**
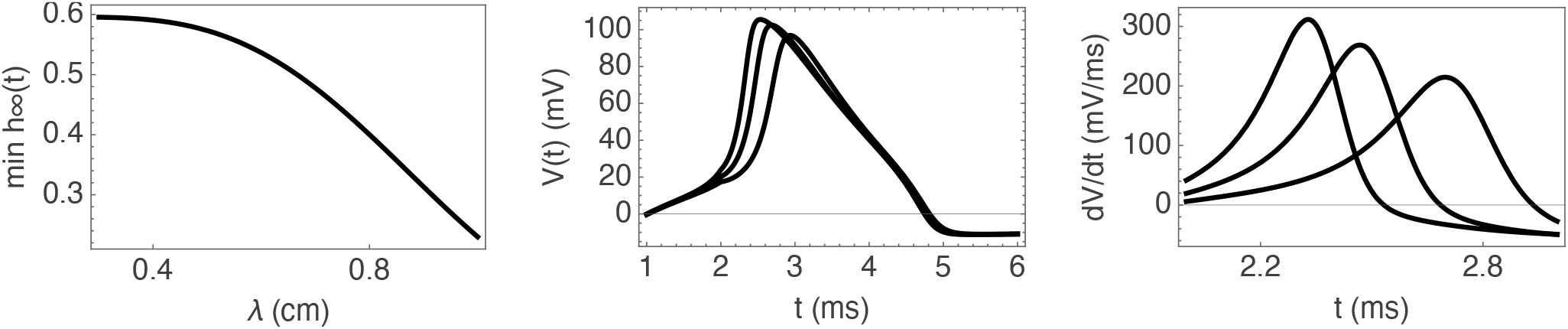
Availability-based realization of an excitability-reserve bias and its spike-shape signature. *Purpose:* a concrete, explicitly computable proxy for a deterministic excitability-reserve component of *η*(Λ) (an *η*_kinetic_-like effect) and an experimentally accessible correlate of reduced regenerative reserve (upstroke slowing). *Scope:* this figure is illustrative and is not used in the bounded-depth derivation; it does not affect the length invariance of Var[ln *v*]. **Left:** Effective minimum Na availability *h*_eff_ (λ) = min_*t*∈[0,*T*]_*h*_*∞*_ (*V*_end_(*t*; λ)), computed from the end voltage *V*_end_(*t*) produced by an illustrative passive end-sensing construction in which a clamped depolarization waveform *V*_*c*_(*t*) approaches a sealed end at speed *v*_app_; increasing λ reduces attenuation and increases *V*_end_(*t*), thereby lowering *h*_eff_ under depolarizing conditions. **Middle:** Single-compartment Hodgkin–Huxley spikes evoked by an identical stimulus current, with all parameters held fixed and only the initial inactivation set to three representative values *h*(0) spanning the range of *h*_eff_ (λ). **Right:** Corresponding upstroke slope traces *dV/dt*, showing a monotone reduction in max(*dV/dt*) with reduced initial availability.

## SUPPLEMENTARY METHODS: MONTE CARLO ILLUSTRATION OF BOUNDED VERSUS UNBOUNDED MULTIPLICATIVE DEPTH

Synthetic slowdown ratios were generated by a log-additive multiplicative process,

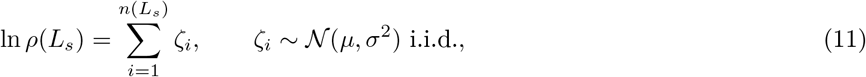

where *L*_*s*_ denotes the total soma-to-terminal branch length (not the distance-from-tip coordinate used elsewhere). Branch lengths *L*_*s*_ were drawn from a truncated log-normal distribution supported on 600–4570 *µ*m, chosen to approximate the empirical length distribution. To define a length-dependent “multiplicative depth” *n*(*L*_*s*_), a coarsegraining length *𝓁* is introduced, that converts a continuous length into an integer step count. Importantly, *𝓁* is a modeling convention (a resolution scale for *n*(*L*_*s*_)), not a claim of a special anatomical segment length. In Fig. 2 *𝓁* = 200 *µ*m is set as a convenient order-of-magnitude discretization.

Code is provided in an accompanying Mathematica notebook:

https://www.wolframcloud.com/obj/marom/Published/SuppFigMarom2026(1DL).nb

The unbounded model assumes the effective depth increases with length,

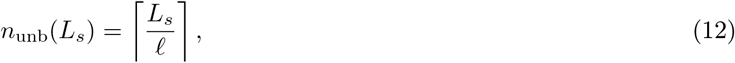

whereas the bounded model caps the depth at a finite saturation value,

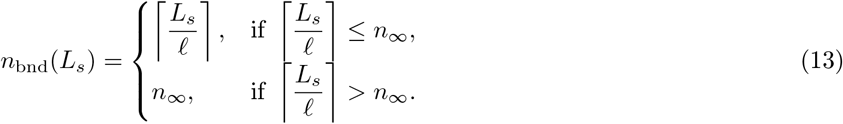

Here *n*_*∞*_ is a *finite asymptotic effective depth* (the maximal number of independent multiplicative factors permitted by the model at large *L*_*s*_), not mathematical infinity. Because *𝓁* is a coarse-graining convention, the associated saturation occurs on the effective length scale set by *n*_*∞*_*𝓁*; this scale is part of the model representation and is not uniquely identified as a physiological length.

Per-step parameters were chosen so that, in the bounded model at saturation (*n*(*L*_*s*_) = *n*_*∞*_), the marginal distribution matches the empirical branch-level fit ln 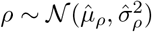. Since at saturation

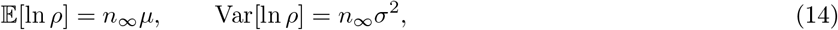

matching gives

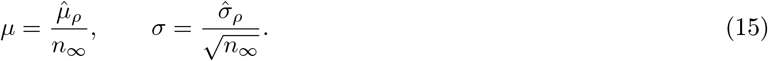

For Fig. 2 (top panels) *n*_*∞*_ = 8 was used as a representative finite cap, yielding *µ* = − 0.04929 and *σ* = 0.09678 from 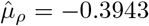 and 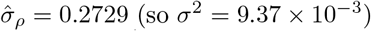 (so *σ*^2^ = 9.37 × 10^−^3).

For each model, *N* = 2 × 10^4^ branches were simulated. Scatter plots show ln *ρ* versus *L*_*s*_ with equal-count binned central trends and marginal histograms pooling ln *ρ* across branches. Bottom panels show equal-count length-bin estimates of 𝔼 [ln *ρ*] and Var[ln *ρ*] compared to the expectations 𝔼 [ln *ρ*(*L*_*s*_)] = *n*(*L*_*s*_)*µ* and Var[ln *ρ*(*L*_*s*_)] = *n*(*L*_*s*_)*σ*^2^. For visual clarity, Fig. 2 displays only *L*_*s*_ ≤ 3.3 mm.

Code is provided in an accompanying Mathematica notebook: https://www.wolframcloud.com/obj/marom/Published/MonteCarlo2r.nb

